# How the visual brain can learn to parse images using a multiscale, incremental grouping process

**DOI:** 10.1101/2024.06.17.599272

**Authors:** Sami Mollard, Sander M. Bohte, Pieter R. Roelfsema

## Abstract

Natural scenes usually contain many objects that need to be segregated from each other and the background. Object-based attention is the process that groups image fragments belonging to the same objects. Curve-tracing tasks provide a special case, testing our ability to group image elements of an elongated curve. In the brain, curve-tracing is associated with the gradual spread of enhanced neuronal activity over the representation of the traced curve. Previous studies demonstrated that the tracing speed is higher if curves are far apart than if they are nearby. One hypothesis is that a larger distance between curves permits activity propagation in higher visual cortex areas. In these higher areas receptive fields are larger and connections exist between neurons representing image regions that are farther apart (Pooresmaeili et al., 2014). We propose a recurrent architecture for the scale-invariant tracing of curves and objects. The architecture is composed of a feedforward pathway that dynamically selects the appropriate scale for tracing, and a recurrent pathway for propagating enhanced neuronal activity through horizontal and feedback connections, enabled by a disinhibitory loop involving VIP and SOM interneurons. We trained the network using a biologically plausible reinforcement learning scheme and observed that training on short curves allowed the networks to generalize to longer curves and 2D-objects. The network chose the scale based on the distance between curves and the width of objects, just as in human psychophysics and the visual cortex of monkeys. The results provide a mechanistic account of the learning and execution of multiscale perceptual grouping in the brain.

**Significance Statement:** In our perception, image elements that belong to the same object are grouped by object-based attention. Object-based attention corresponds to an enhanced neuronal representation of the image elements that are grouped in perception, in multiple areas of the visual cortex. During perceptual grouping tasks, this enhanced neuronal activity spreads gradually over an object representation, with a speed that depends on the distance between the relevant object and other objects. Here we propose a neuronal mechanism that learns scale-invariant spread of object-based attention and accounts for psychophysical observations in human observers and the pattern of neuronal activity in the visual cortex of monkeys. This work sheds light on the mechanisms for multiscale object-based attention in the visual cortex.

## Introduction

Natural scenes usually contain several objects that need to be segregated from each other and from the background to guide behavior. Previous studies demonstrated that neurons in areas of the visual cortex that encode the elements of a relevant object incrementally enhance their firing rate during perceptual grouping tasks (1). This neuronal process corresponds to the gradual spread of “object-based attention” in perceptual psychology (2, 3). The rules determining what groups with what were already studied in the first half of the 20th century by the Gestalt psychologists (4, 5). One of their rules is connectedness, because image elements that are connected to each other tend to belong to the same object and group in our perception. Another rule is that of good continuation, describing that collinear contours usually belong to the same object.

Early theories of perception suggested that Gestalt rules are applied pre-attentively and in parallel across the visual field (6), and indeed, some basic forms of grouping related to, for example, the perception of shape take place in parallel across the visual field (2, 3). However, grouping is flexible, because we can also group the features of objects that we never saw before. These groupings appear to form incrementally if image elements are grouped indirectly, via other image elements. On example of such seriality occurs in curve tracing tasks, in which participants report if two elements belong to the same curve (Fig. 1A). Jolicoeur et al. (7) showed that the reaction time is proportional to the length of the curve that subjects have to trace. This serial process is also reflected by neurons in the visual cortex. Neurons in the visual cortex of monkeys (1) and humans (8) with a receptive field (RF) on the target curve increase their firing rate relative to neurons with RFs on a distractor curve (Fig. 1A,B). This response enhancement does not occur during the initial feedforward visual response, triggered by the onset of a visual stimulus in the RF, but after a delay. The latency of the response enhancement depends on the distance between the start of the tracing process and the position of the RF. It occurs later for neurons with RFs farther along the curve (9), in accordance with the gradual spread of enhanced activity over the representation of the target curve (Fig. 1A). The spread of an enhanced response corresponds to the gradual spread of object-based attention across the relevant curve (10, 11).

**Figure 1.**
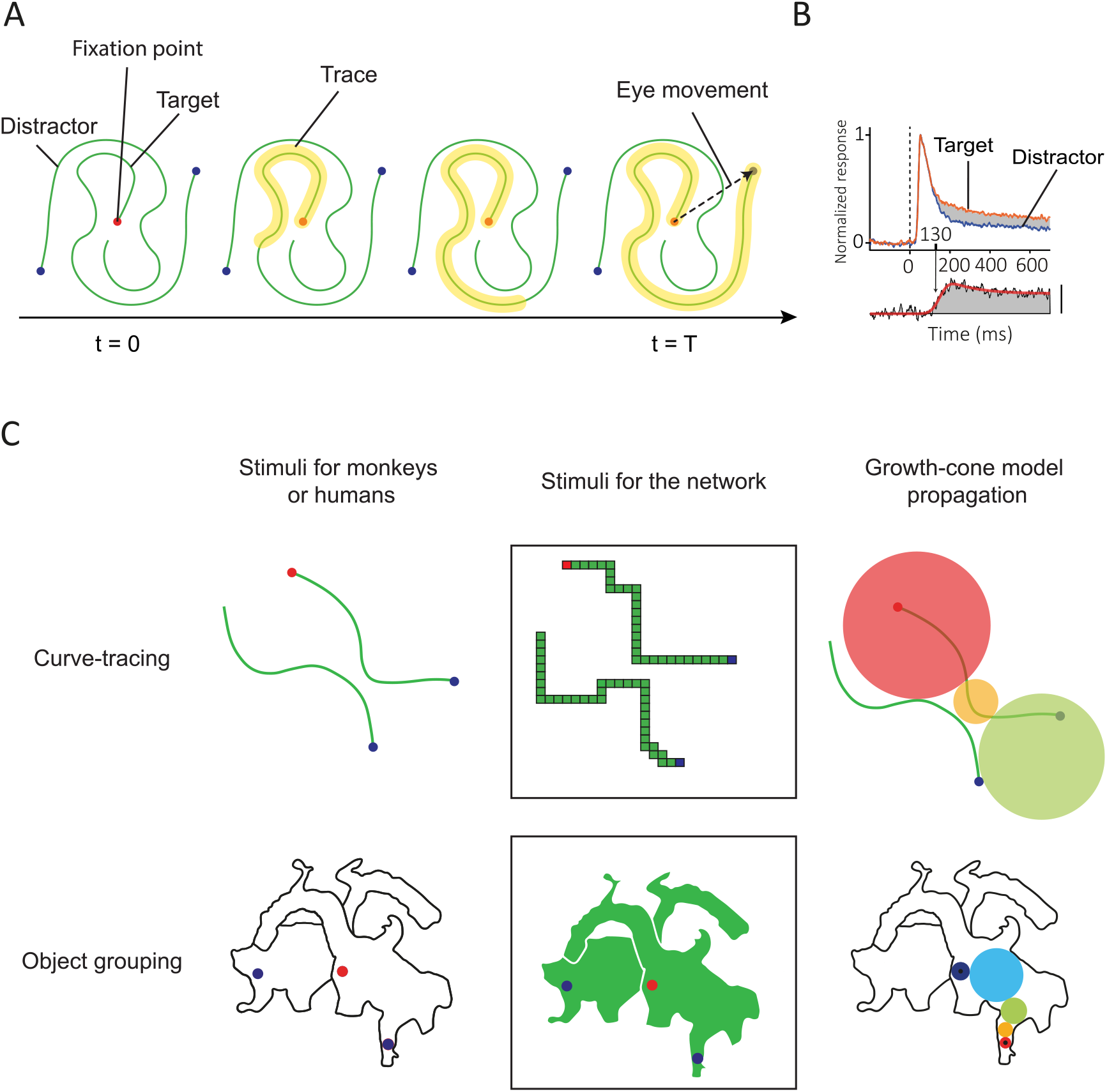
Tasks used to probe the dynamics of object-based attention. **A**. In this example curve tracing task, the subject must make an eye movement toward the blue dot connected to the red fixation point. Extra neuronal activity spreads across the representation of the target curve in the visual cortex (yellow). **B**. Recordings in the visual cortex of monkeys revealed that the target curve is labeled with enhanced neuronal activity (orange) compared to the distractor curve (blue). **C**. Curve tracing and object grouping. In the curve tracing task, a growth-cone model of attention can account for the time-course of neuronal responses, proposing that enhanced activity spreads at multiple scales. If curves are far apart, the model uses large RFs (large circles) and smaller RFs when curves are nearby. The object grouping task is analogous to the curve-tracing task but takes place with two-dimensional objects. Subjects determine which dots are on the same image component.

The curve-tracing speed depends on the distance between the target curve and the distractor curves and the curvature. Tracing slows down if distractors are nearby (9, 12) or if the traced curve has a high curvature (13). Pooresmaeili et al. (9) tested the influence of the distance between RFs on tracing speed in the visual cortex of monkeys. In their experiment there was a bottleneck where the target curve came close to the distractor (Fig. 1C, top). Narrowing of the gap between the curves delayed the onset of the response enhancement, but only for parts of the target curve that were behind the bottleneck. A possible explanation of the influence of distance between curves is that the spread of enhanced neuronal activity across the target curve occurs in multiple cortical areas in which the size of RFs differ (9). When the target curve is close to a distractor, the propagation of the enhanced activity would occur in low-level areas, such as the primary visual cortex, where RFs are small so that they do not fall on both curves. In these low-level areas, connections between neighboring neurons interconnect neurons with nearby RFs so that tracing speed is low. When the curves are farther apart, the spreading could occur in higher areas where the neurons have larger RFs and connections bridge across larger distances in the visual field. A “growth-cone” model for the spread of object-based attention proposed that fastest progress is made in the area where RFs falling on the target curve almost touch the distractor curve (Fig. 1C). An interesting implication of the growth cone model is that the reaction time in curve-tracing only depends little on the viewing distance from the display. When the subject approaches the stimulus, the length of the curves, measured in degrees of visual angle, increases. However, the distance between curves also increases and grouping can rely on neurons in higher visual areas with larger RFs. These two effects cancel each other and the reaction time remains the same (14).

Jeurissen et al. (15) extended the growth cone model to 2-D images. In their study, human participants reported whether two dots were on the same object or not. The pattern of reaction times was best explained by a version of the growth-cone model, in which image regions are incrementally labeled with enhanced activity. The enhanced activity started at one of the dots and propagated across the target surface through neurons with RFs with sizes that were small enough to stay within the object boundaries (Fig. 1C, bottom).

In a previous study, Marić & Domijan (16) proposed an instantiation of the growth-cone model for curve-tracing in a neural network with multiple scales. They used hard-coded, multi-scale Gabor filters and manually selected the synaptic weights to select the appropriate scale. Their network qualitatively reproduced the results observed in monkey visual cortex, but the model did not yet generalize to 2-D images.

Here we examine how recurrent neural networks with units with several RF sizes learn to trace curves and to fill in objects in a scale-invariant manner. Building on prior work (16, 17) we developed an architecture with segregated populations of units that either represent the stimulus veridically because of pure feedforward connectivity or can be modulated through horizontal and feedback connections. A key innovation is the inclusion of a trainable, biologically inspired disinhibitory loop that enables attentional modulation (18). Furthermore, we implement a biologically plausible learning rule that can train networks by trial-and-error. The only feedback that the model receives is a reward if it makes the correct choice, like how monkeys are trained on a curve-tracing and region-filling tasks. Specifically, we used RELEARNN (17, 19) to determine how synapses modify their connection strength, using information both local in space and time.

Our approach addresses (1) how the networks learn to group image elements of the same objects, (2) the mechanisms allowing networks to learn to propagate enhanced activity at multiple spatial scales. We report that training on short curves allowed the model to generalize to long curves and to 2-D shapes. Furthermore, the networks predicted human reaction times during image parsing tasks and the spread of neuronal activity in the visual cortex of monkeys.

## Model and task

We trained the networks on a curve-tracing task that has been used during electrophysiological recordings in the visual cortex of monkeys (9) (Fig. 1A). The monkeys first had to direct their gaze to the fixation point before the stimulus appeared, but in the version of the task for the network, the entire stimulus was presented at once. We also tested how well models proficient in curve-tracing generalize to the parsing of 2-D image regions (15, 20). The task of the model was to select a blue pixel on a curve or object that was cued with a red pixel (Fig. 1C) as target for an eye movement.

As in previous work (17), we included a feedforward and recurrent processing group of units (Fig. 2A). Neurons in the feedforward group only receive feedforward input and propagate the information to higher layers. They represent the stimulus veridically and are responsible for the selection of the appropriate scale. Neurons in the recurrent group receive feedforward connections from lower layers, feedback from higher layers and horizontal connections from neighboring units in the same layer. They are responsible for the propagation of enhanced neuronal activity. The presence of some units that propagate the enhanced activity and others that do not is in accordance with neurophysiological results (21–24). Neurons that engage in incremental grouping are enriched in layers 2, 3 and 5 of the visual cortex, whereas neurons that veridically represent the feedforward input are mostly situated in layers 4 and 6 (21–23). In the model, units in the feedforward group gated the units of the recurrent group, so that they could not spread enhanced activity if the corresponding feedforward unit was not active (Fig. 2A). This feature prevented the spreading of enhanced activity into image regions not occupied by curves or objects.

**Figure 2.**
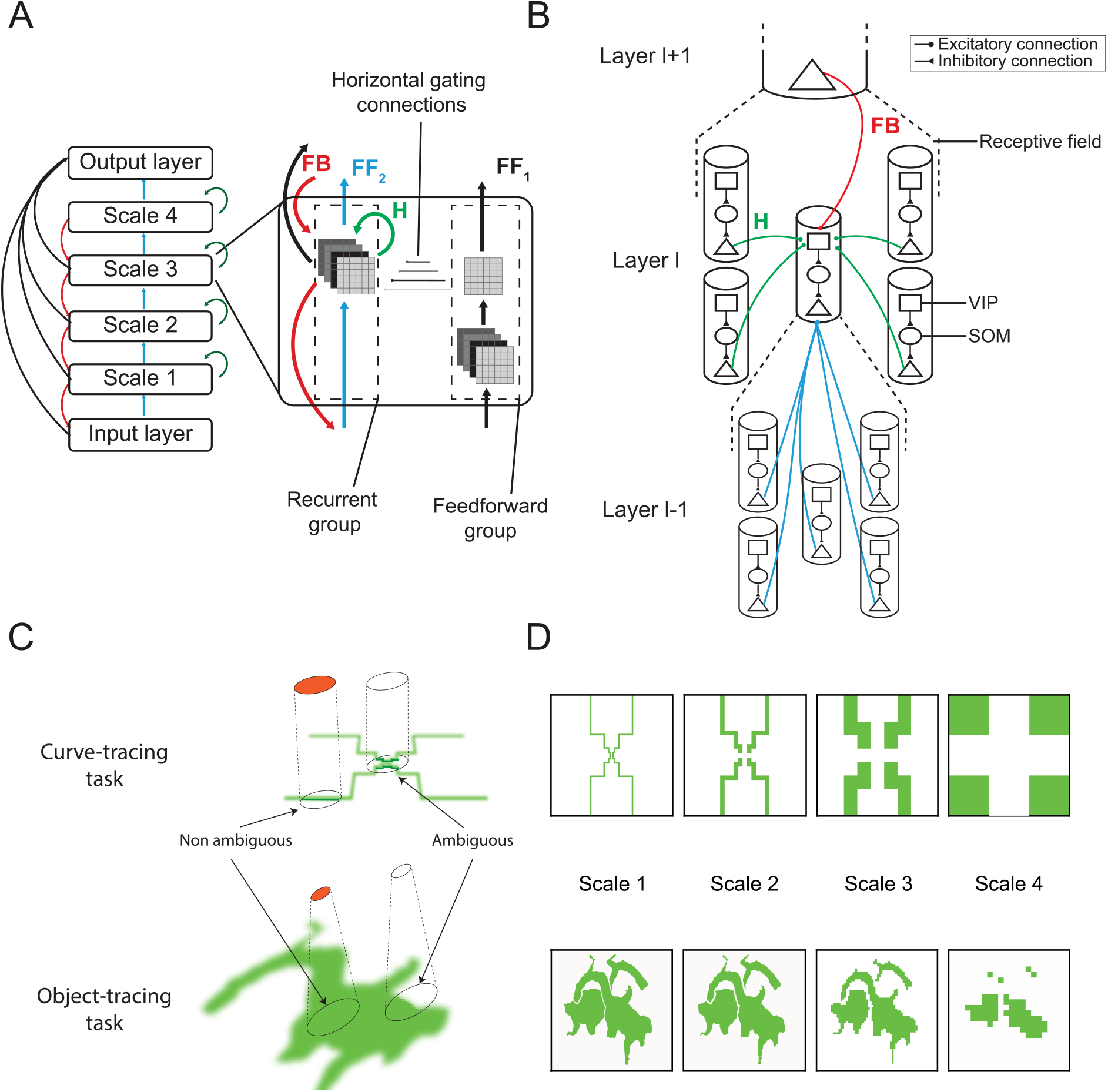
Model. **A.** The network incorporated four spatial scales. Units in the feedforward network are responsible for scale selection, whereas units in the recurrent network spread enhanced activity. This spread is gated by feedforward units with overlapping RFs. **B**. Disinhibitory interactions between recurrent units. Pyramidal neurons receive feedforward from pyramidal units in the layer below. They also receive top-down input from units in the layer above and horizontal input from their neighbors through a disinhibitory loop composed of VIP and SOM interneurons. **C**. Scale selection in the feedforward network. In the curve tracing task, units in the feedforward group are only active (orange) if the image elements in the RF are connected and colinear. The cells are not active otherwise so that the enhanced activity cannot spill from the target to the distractor curve. In the object-parsing task, feedforward units are active if their RF falls on a homogeneous image region. **D**. Activity in the feedforward network in the curve tracing (top) and the object parsing task (bottom). The representation in higher layers is coarser, but RFs are larger and propagation can proceed faster.

### Feedforward network

The neural network has four spatial scales, and the feedforward units are responsible for scale selection. They are trained to only activate if the stimulus in their RF unambiguously belongs to a single object (see below) to prevent “spilling” of enhanced activity from the target to the distractor object in the recurrent network. In the curve-tracing task, the stimulus in an RF is unambiguous if all pixels are connected to each other and colinear (Fig. 2C). Hence, the model selects a finer spatial scale for propagation if the target and distractor curves are nearby, or if the target curve has a high curvature (13). In the object-parsing task, the stimulus inside a receptive field is unambiguous if it doesn’t contain a boundary (Fig. 2C). Upon presentation of the stimulus, feedforward units propagate activity to higher layers if the stimulus in their RF is unambiguous, resulting in a multiscale base-representation (Fig. 2D). The activity *X*^*l*^ of units in the feedforward group in layer *l t* is determined by:

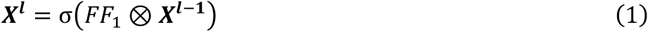

*FF*_*1*_ is the set of feedforward connections in the feedforward network, σ is a ReLU activation function and ⊗ indicates convolution.

### Recurrent network

Units of the recurrent network learned to spread enhanced neuronal activity over the representation of the target object, to label it as one coherent perceptual group. They are connected by feedforward, horizontal and top-down connections. The propagation of enhanced neuronal activity relied on disinhibition, which was advantageous because we used the RELEARNN learning rule (see below) (19) requiring the network to reach a stable state, which is not guaranteed with ReLU nonlinearities (Material and Methods). The disinhibitory connection scheme (Fig. 2B) was inspired by the interactions between inhibitory neurons in the mouse visual cortex during figure-ground segregation (18) and has been used in previous models of attentional selection (25, 26). It ensured stable and expressive networks that learned to trace curves with an arbitrary length.

Specifically, the feedback and horizontal connections activated vasoactive intestinal peptide-expressing (VIP) interneurons. The VIP neurons inhibit somatostatin-expressing (SOM) interneurons, which inhibit pyramidal neurons (18, 27). Hence, VIP neurons disinhibited the pyramidal neurons. During curve-tracing, VIP neurons incrementally disinhibit pyramidal units that represent the target object, which is thereby labeled with enhanced activity. The advantage of the disinhibitory scheme is that activity of pyramidal neurons cannot be higher than what it would be without SOM inhibition, preventing run-away excitation that can occur in recurrent networks with excitatory units that directly excite each other.

The activity *Y*^*l*^(*t*) of units in the recurrent group in layer *l* and time *t* is determined by:

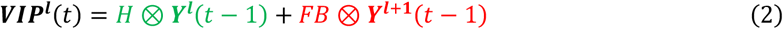

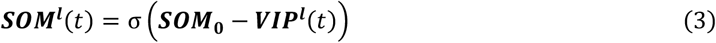

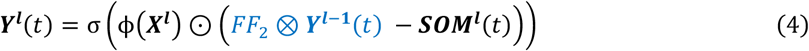

FF_2_ and FB represent feedforward and feedback weights between the layers of the recurrent network, respectively, and H the horizontal weights within a layer (colors correspond to those in Fig. 2A,B). SOM_0_ is the default activation of SOM neurons in the absence of feedback, σ is a ReLU activation function and ϕ a gating function defined by:

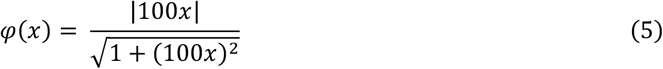

This gating function ensures that propagation of the enhanced activity can only occur if the feedforward units ***X***^***l***^ are active. ⊙ represents the element-wise product between recurrent and feedforward units.

### Output layer

The output layer was retinotopically organized and units in the output layer learned to represent the Q-value (28), which is the expected reward for the selection of an eye-movement to one of the pixels. It was connected to all layers, including the input layer with skip connections, allowing it to integrate the progress in incremental grouping across all scales.

The estimated Q-value was determined when the network reached a stable state at time T:

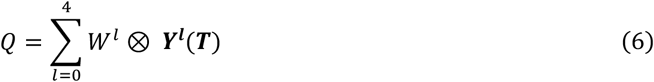

During training, the model chose to make an eye-movement toward the position with the highest Q-value with probability 1 − *ε*. With probability *ε* other actions were explored, by sampling a random action a from the Boltzmann distribution *P*_*B*_ :

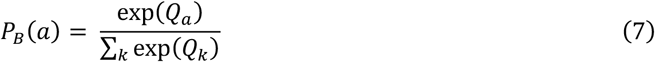

We simulated the selection of an eye movement but not the eye-movement itself or the shift of the visual image caused by it.

## Training

We trained the networks in two steps. First, we trained feedforward networks, one for the curve-tracing task and one for the object-parsing task. The feedforward units for curve-tracing were trained to activate if all the image elements inside their RF were colinear and connected (Fig. 2C,D) using backpropagation, using binary cross entropy loss. We used the Adam optimizer (29) and a learning rate of 10^−3^. Similarly, the feedforward units for object-parsing were trained to activate if all input units in their RF were active (Fig. 2C,D). Each scale was trained independently on the same set of stimuli. The stimuli were random curves on a 36×36 grid for curve tracing, and random objects on a 72×72 grid for object-parsing. We assumed that the tuning of feedforward units had emerged during visual experience (30–34), prior to training on curve-tracing or object-parsing. We note, however, that the non-biological backpropagation learning rule could be replaced by biologically plausible reinforcement learning rule (35). The weights in the feedforward group were frozen during training of the recurrent weights.

Weights in the recurrent network were updated using RELEARNN, a local learning rule inspired by the Almeida-Pineda algorithm (2, 17, 19, 36) that uses three phases. First, the activity of input neurons and units of the feedforward network determines the propagation of activity until convergence to a stable state. We considered that a stable state was reached if the activity of neurons between two consecutive timesteps was the same or after 30 timesteps. The network then selected an eye movement, as described above, and an attention signal originating from the winning action propagated through an accessory network to determine the influence of each neuron on the selected action (for details see ref. (17)). The network received a reward r of 1 unit in case the eye movement was correct and 0 otherwise. It then computes a reward prediction error *δ* :

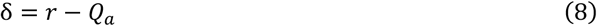

Where Q_a_ is the activity of the selected output unit. The reward-prediction error *δ* could be broadcasted to the whole network by a neuromodulatory signal (37) and become available at all synapses. The reward-prediction error *δ* and the attentional feedback signal are available at the relevant synapses and determine the weight update, together with the pre- and post-synaptic activity. Hence, the required information is available locally at the synapse, making RELEARNN biologically plausible. We used a curriculum for training (17), a strategy also used to train monkeys. The network was first presented with curves of 3 pixels. Once the network achieved 85% accuracy during a test phase with fixed weights and no exploration, we added one pixel to the curve and repeated this procedure until the curves were 7 pixels long.

To reduce the computation time, we used weight sharing, although this is biologically implausible. In previous work, we showed that networks with a similar structure learned to trace curves with or without weight-sharing, but that the number of trials needed to learn the task without weight-sharing was ∼7 times larger (17). Hence, our results are likely to generalize to learning rules without weight sharing.

## Results

### Curve-tracing task

We trained 5 networks on a curve-tracing task, which was a variant of a task that has been used for electrophysiological recordings in monkeys visual cortex (9) (Fig. 1A,C). In the current version of the task, the stimulus (illustrated in Fig. 1C) was presented at once (Fig. 3A), and the network was rewarded if it selected an eye movement to a blue pixel that was connected by a curve to a red pixel. We trained the networks with a curriculum in which the length of the curves was increased until they were 7 pixels long. All networks converged (criterion of 85% correct) within an average of 23,200 trials. To examine whether the networks memorized specific curve configurations (38) or learned a general grouping rule we generated fifteen curves with a length of 30 pixels. The accuracy of the networks was 100%, confirming that they learned a general solution. Units of the recurrent network had learned to iteratively spread enhanced activity across the representation of the target curve.

**Figure 3.**
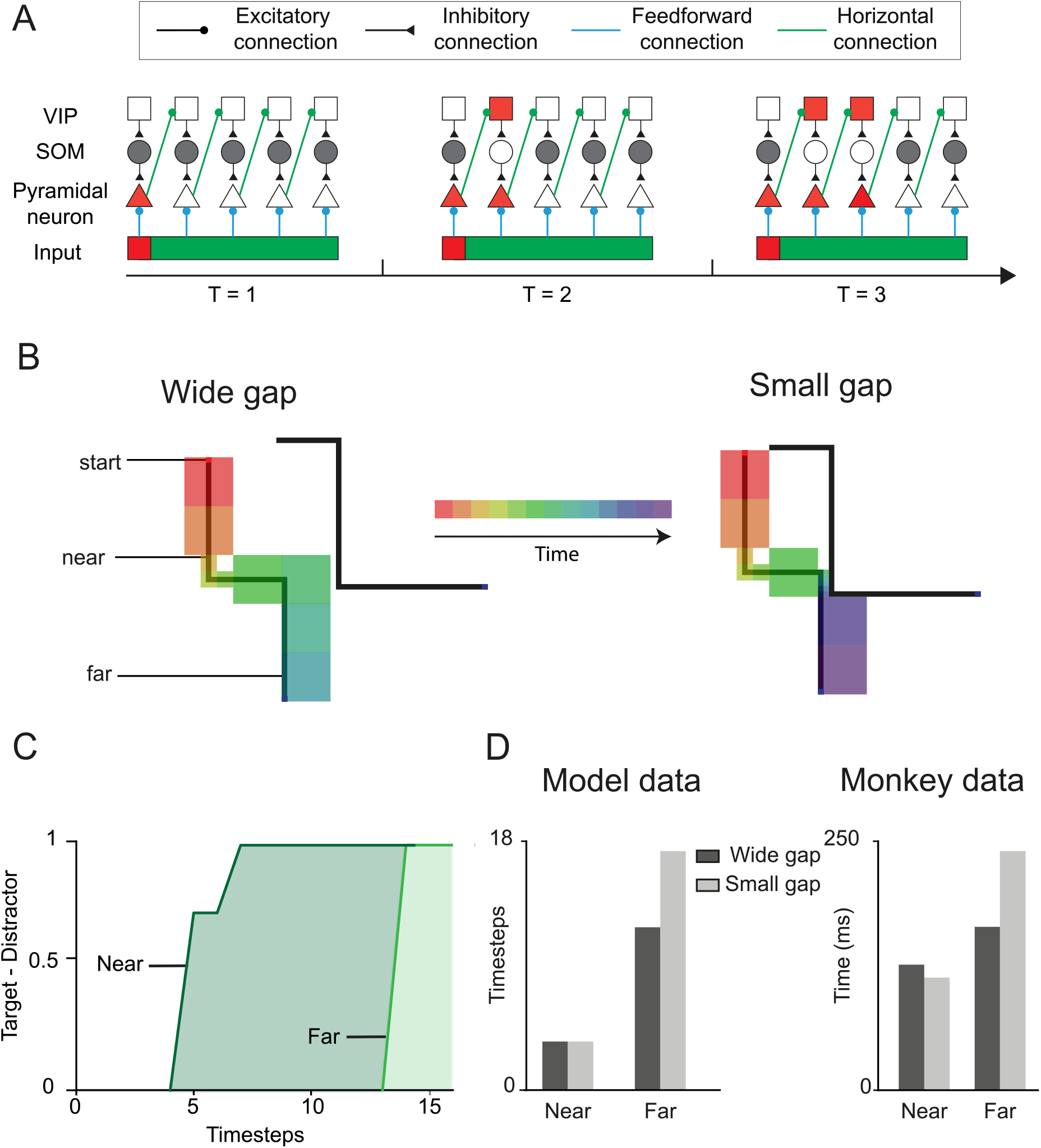
Networks propagate the response enhancement at multiple scales. **A**. Spreading of activity by disinhibition. At t=1 the visual stimulus is presented, and the red pixel activates a pyramidal unit, which in turn activates VIP units in an adjacent column through horizontal connections (green), which cancels SOM inhibition and activates the next pyramidal cell at t=2. The disinhibition then propagates further along the target curve. **B**. Stimuli with a larger or narrow gap, which were not used in training. The size of the squares represents the scale that was used by the networks and color represents the time at which the attentional tag reached the RF. When the curves are far apart, the tracing process is faster because it occurs at a larger scale. **C**. We measured the activity of units with RFs before or after the gap. The activity elicited by the target curve was higher, and the response enhancement occurred later for units with RFs behind the gap (far) than for units with RFs before the gap (near). **D**. Latency of the response enhancement in the model (left) and in area V1 of monkeys (right). The response enhancement did not depend on gap size for near RFs before the gap. A narrow gap delayed the response enhancement of units with far RFs behind the gap.

The model learned to use the disinhibitory circuit for spreading enhanced activity through the recurrent network (Fig. 3A). The model’s SOM interneurons are spontaneously active and inhibit the pyramidal neurons with overlapping RFs and the feedforward input does not overcome this inhibition. Through trial-and-error learning, the network learns that the target curve begins with a red pixel. Specifically, learning increases the feedforward weights from the red pixel start so that the units that represent this location overcome SOM inhibition (T=1 on Fig. 3A). This activity activated adjacent VIP interneurons via horizontal and feedback connections, which inhibited the adjacent SOM interneurons and disinhibited the pyramidal cell. These interactions repeated, causing the propagation of disinhibition across the target curve.

We next examined how the model’s tracing speed depends on the distance between the target and distractor curves. We presented stimuli with a bottleneck where the target curve came close to the distractor (9) (Fig. 3B) and examined the activity of units with RFs before and after the bottleneck. The latency of the response enhancement, measured as the time-step where activity reached 90% of its maximum, was shorter for units with RFs before the bottleneck than for units with RFs behind (Fig. 3C). Narrowing the gap between the curves delayed the onset of the response enhancement, but only for RFs after the bottleneck (Fig. 3D), just as has been observed in the visual cortex of monkeys (9). We next examined the scales that were selected by the model (Fig. 3B) and noticed that the small gap enforced the spread of enhanced activity at lower network levels, where RFs were smaller. The propagation speed was also influenced by curvature because the network resorted to spreading activity at lower network levels if image elements in the larger RFs were not colinear. This results is in agreement with experimental evidence in humans that a higher curvature decrease tracing speed (13).

### Object-parsing task

We next probed the ability of the networks to group image elements of 2-D objects, using a variant of an object-parsing task used in human participants (15, 20, 39). The participants reported whether a cue feel on the same object as the fixation point. The version for the network was like the curve tracing task of the previous section. The fixation point was a red pixel, and we placed a blue pixel on the same 2-D object and another one on a second object. The network had to plan an eye movement to the blue pixel on the same object as the fixation point (Fig. 4A).

**Figure 4.**
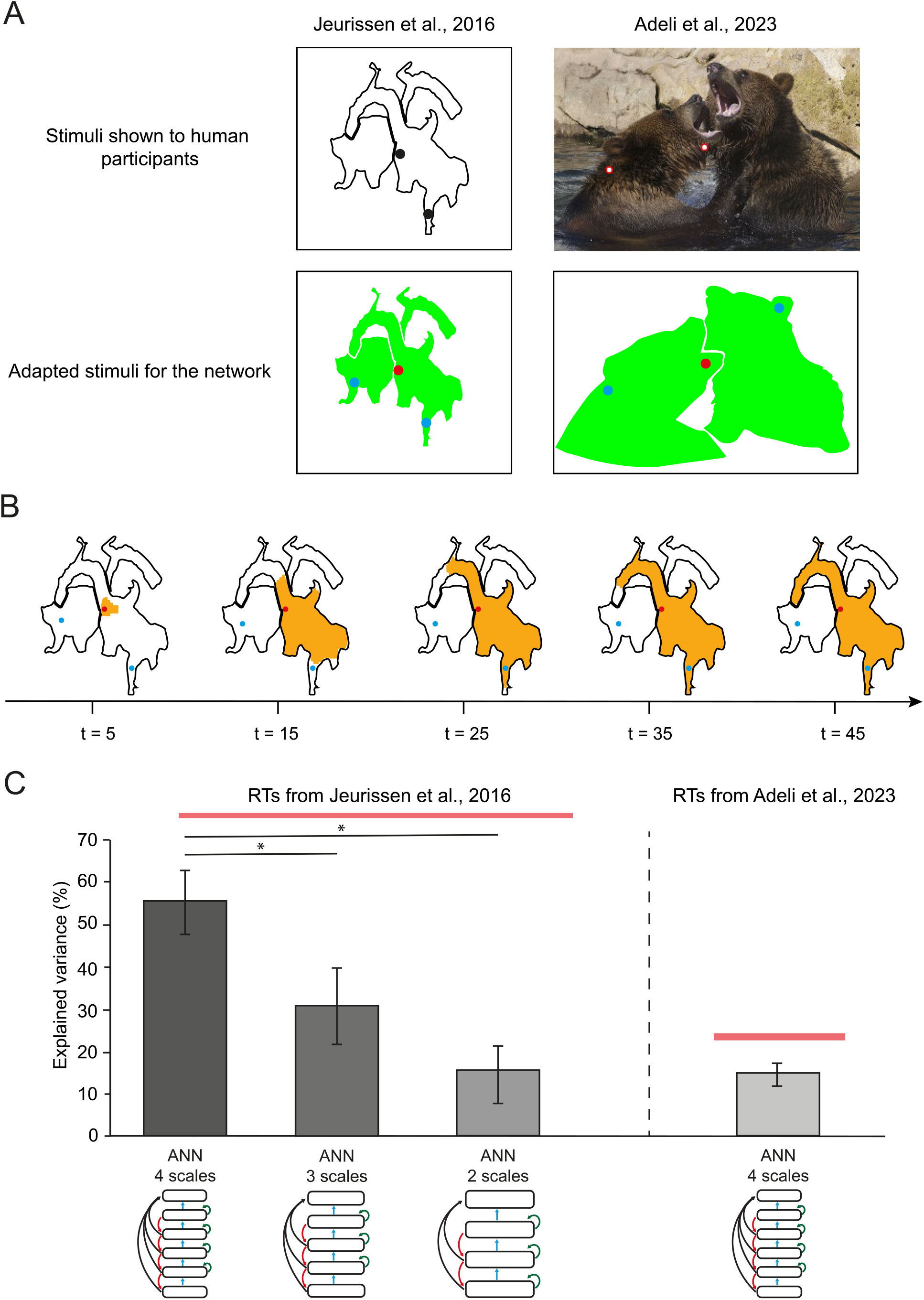
Object-tracing task. **A.** We examined how well the model could explain human reactions times (RTs) in image parsing tasks. We presented simplified images where objects were filled homogeneously and background locations were empty. **B**. Illustration of the spread of enhanced activity (orange) starting from the red dot. Large homogeneous image regions are filled faster (steps 5-15) than narrower regions (steps 25-35). **C**. Explained variance in human reaction times collected by Jeurissen et al. (10) and Adeli et al. (19). We fitted the reactions times with network models with either 2, 3 or 4 scales. Note that the quality of the fit improves for more scales. Error bars, standard errors. Red line, noise ceiling. *, P<0.05.

The models had the same architecture as those for the curve-tracing task (Fig. 2). We started with the recurrent units that had been trained on the curve-tracing task and retrained their weights for the object-parsing task using the same RL process. For the feedforward groups, we used networks that were specifically trained for scale selection in the object-parsing task. All 5 networks reached an accuracy higher than 85% within 100 trials when they were tested in the object-parsing task. This additional training did not influence accuracy in the curve-tracing task, which remained 100%, implying that the recurrent architecture can account for both curve-tracing and object-parsing. The networks parsed the object that was cued by the red fixation point by spreading enhanced activity over its representation (Fig. 4B), just as has been observed in V1 of humans with fMRI (40).

We next investigated the dynamics of the parsing process by comparing the processing time in the model to human reaction times in two studies. The first experiment was by Jeurissen et al. (15) and presented scrambled shapes, which could not be recognized by the participants (Fig. 4A). To model the reaction time, we estimated the timestep at which units in the output layer whose receptive field fell on the blue pixel enhanced their response by 90%. The model explained 55% of the variance of human RTs. The heuristic growth-cone model (Fig. 1) accounted for 63% of the variance, which is close to the noise ceiling of 67%. A key difference between the growth-cone model and its implementation as neural network is the number of scales. The neural network used four scales, whereas the number of scales used by the growth-cone model was not bounded. To further examine the dependence on the number of scales, we tested models with fewer scales. We observed that the models with 2 scales and 3 scales explained significantly less variance than the model with 4 scales (p < 10^−3^ and p = 0.03, respectively, see Methods) (Fig. 4C).

We replicated these findings modeling data from a second study that presented naturalistic images from the COCO dataset (41). We simplified the task by presenting object masks to the neural network and we measured the number of timesteps before the enhanced response reached the blue pixel on the same object as the fixation point (Fig. 4A). The neural network accounted for 15% of the variance in the human reaction times, which is close to the 16% accounted for by the growth cone model and to the relatively low noise ceiling of this data set of 24%. These results imply that that the neural network accounted well for the patterns of human reaction times in image parsing tasks.

## Discussion

In this study, we used neural networks to understand the propagation of object-based attention in curve-tracing and object-parsing tasks and how it can be learned (42). The processing time in curve tracing tasks increases linearly with the length of the curve that needs to be traced. In the visual cortex of monkeys (1, 9) and humans (8, 40), curve-tracing is associated with the gradual propagation of enhanced neuronal activity along the representation of the target curve (8). The speed of the tracing process depends on the curvature of the target curve and on its distance to distractor curves (9, 13). Recent studies started to examine the neuronal processes responsible for the parsing of 2-D surfaces and natural objects where the processing time also depends on the distance between the image locations that need to be grouped, and on the width of the image regions connecting them (15, 20).

We trained a recurrent neural network to perform a curve-tracing task using trial-and-error learning, mimicking how monkeys are trained. With minimal additional training, the same networks could also parse spatially extended objects. Remarkably, the model accounted for many neurophysiological and psychophysical findings. We reproduced the pattern of reaction times in humans in curve-tracing and image-parsing tasks, including the influence of the curvature and distance between curves and the width of image regions. Furthermore, the model developed a strategy to incrementally label the relevant curve or image region with enhanced neuronal activity, just as is observed in the visual cortex. The model thereby provides a neural network implementation of the conceptual “growth-cone” model (Fig. 1C) (9, 13, 15), explaining why grouping speed depends on the distance between curves and the width of object regions. With only 4 scales, the network’s predictive power approached that of the growth-cone model, which had not yet been implemented as neural network and did not commit to a specific number of scales.

The model thereby goes beyond previous work on incremental grouping by neural networks. It learned to trace curves by trial-and-error at multiple scales, whereas previous networks for curve-tracing used a predetermined connectivity scheme and did not generalize to the parsing of 2D image regions (16, 43). The model also extends studies (44, 45) that introduced a specialized hGRU unit for horizonal interactions to solve image parsing tasks. These models computed a measure of uncertainty, which had to be transformed into a measure of human reaction time and they accounted for less variance than the present model (21% vs. 55% for Jeurissen et al. (15), and 7% vs. 15% for Adeli et al. (41)).

The new model incorporated several key features, inspired by the neurophysiology of the visual cortex. Firstly, we created a hierarchically organized network with larger RFs in higher layers, which enabled the faster grouping of straight curves and homogeneous image regions (16). Secondly, we used a feedforward and a recurrent network with separate roles. Units of the feedforward network provided a veridical representation of the stimulus, providing a stable scaffold for the spreading of enhanced activity in the recurrent network. There are neurons with similar properties in input layers 4 and 6 of the visual cortex of monkeys (21–23), which do not participate in incremental grouping. We trained the feedforward units to only respond to image regions that unambiguously belonged to a single object, enabling the network to select the appropriate scale. The feedforward network gated the recurrent units and thereby enabled a set of recurrent interactions that was appropriate for the scale of the stimulus in the RF (46). Neurons in the recurrent network learned to spread enhanced activity over the representation of the target object. In the cortex, such neurons that participate in incremental grouping are prominent in layers 2, 3 and 5 (21, 23). Thirdly, we implemented a disinhibitory scheme in which VIP interneurons inhibited SOM interneurons, thereby disinhibiting pyramidal units. This implementation is in accordance with experimental evidence for the role of this disinhibitory circuit in figure-ground segregation (18). The disinhibitory circuit has computational advantages, because the maximal activity of the excitatory units in the circuit is bounded by their feedforward input, preventing run-away excitation and guaranteeing that the network reaches a stable state. Furthermore, the disinhibitory circuit enabled the network to trace long curves without attenuation of the response enhancement, unlike in previous work (17).

The present study may inspire future work on the neuronal mechanisms underlying image parsing. For example, one limitation of the present approach is that we pretrained feedforward units at higher network levels to detect stimulus configurations that permit grouping at larger spatial scales, before we used reinforcement learning to train the recurrent network to trace curves. Humans learn to detect colinear configurations during a visual development phase, which extends into late childhood (47). Other perceptual grouping cues, like similarity of luminance, are also learned during development. Future studies could develop biologically realistic learning rules that permit the simultaneous learning of these grouping heuristics at higher network levels and the propagation of enhanced neuronal activity for incremental grouping.

It would also be of interest to generalize the present approach to segment natural images, which is a process that depends on object-recognition (48). Indeed, humans parse upright images more efficiently than images that are presented upside down (39, 49), illustrating how object recognition aids in image parsing. Although the segmentation of natural images is a largely solved problem for transformer-based deep neural networks, the mechanisms for image parsing in the human brain remain only partially understood. The present approach could be generalized to model the recognition of objects in cortical areas, which provide feedback to label individual object parts and low-level features, represented in lower visual cortical areas with enhanced neuronal activity.

In conclusion, the present results provide insight into how brain-like networks learn to integrate grouping cues represented at lower and higher network levels by the spread of enhanced neuronal activity. At a psychological level of description, this process maps onto the spread of object-based attention across all features that are integrated in coherent object representations (10, 11). We look forward to future work, leveraging the highly productive convergence between machine learning, neuroscience and perceptual psychology, to help us better understand how rich, multi-feature object representations emerge in our conscious perception.

## Materials and Methods

### Visual stimuli

We probed the capacity of the network to select the blue pixel that was on the same curve or object as the red pixel. We generally used images with 108×108 pixels with three color channels; red, blue and green (Fig. 1C). To test scale selection, we also presented images with 144×144 pixels for the curve tracing task, and 594×594 pixels for the object parsing task. The larger images did not require fine-tuning because we used weight-sharing.

### Alternatives for the disinhibitory connection scheme

We compared networks with disinhibition to networks in which the excitatory units directly activated adjacent excitatory units. These networks were unstable if we used ReLU activation functions, whose activity is unbounded. When we replaced ReLUs with a squashing non-linearity, the strength of the response enhancement decreased with the length of the target curve so that the network did not learn to trace curves longer than 12 pixels, in accordance with previous work (17). The disinhibitory circuit enabled a stable solution that generalized to longer curves. The network learned a strategy in which pyramidal units in the recurrent network were either inactive because they were inhibited by SOM interneurons or active when the disinhibitory signal reached them. The bi-stability prevented the attenuation of the level of enhanced activity across the curve, allowing the network to trace curves of any length.

### Estimation of human reaction times

To analyze the pattern of human reaction times, we followed the procedures of the studies where they were gathered. To model the results of Jeurissen et al. (15), we analyzed RTs on correct trials with the fixation point and cue on the same object. We removed outliers that deviated from the mean of the 1/RT distribution by more than 2.5 standard deviations. We averaged the RTs across participants, so that there was one RT for every combination of image (20 images) and cue (3 cue positions per image). For Adeli et al. (20) we also analyzed RTs on correct trials with the fixation point and cue on the same object. We computed the average RTs across participants for every combination of image (255 images) and cue (2 cue positions per image). We performed a regression analysis to test how well RTs were predicted by the growth-cone model, and the artificial neural network.

We computed the standard error for the coefficients of determination (R^2^) using Cohen et al. (2003) (50) formula (p. 88):

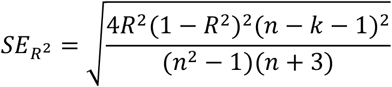

Where n is the number of observations and k is the number of independent variables.

To compare R^2^values between the model with 4 scales and models with 2 or 3 scales, we computed the standard error of the difference:

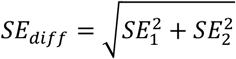

Then, we calculated the Z-score:

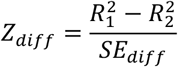

Finally, we determined the p-value using:

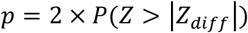

## Acknowledgments

This research has received funding from the European Union’s Horizon 2020 Framework Programme for Research and Innovation under the Specific Grant Agreement No. 945539 (Human Brain Project SGA3, Task 3.7, P.R.R, S.M.B.), Horizon Europe (ERC advanced grant 101052963 “NUMEROUS”, P.R.R.), NWO (Crossover grant 17619 “INTENSE” and NWO-OCENW.KLEIN.178, S.M.B.), “DBI2”, a Gravitation program of the Dutch Ministry of Science (S.M.B.), and Agence Nationale de la Recherche (AN) within Programme d’investissement d’avenir, Institut Hospital Universitaire EQReSIGHIT (ANR-18-590 IAHU-0001, P.R.R.).

We acknowledge the use of Fenix Infrastructure resources, which are partially funded from the European Union’s Horizon 2020 research and innovation programme through the ICEI project under the grant agreement No. 800858.

## References

1. P. R. Roelfsema, V. A. F. Lamme, H. Spekreijse, Object-based attention in the primary visual cortex of the macaque monkey. Nature 395, 376–381 (1998).

2. P. R. Roelfsema, Cortical algorithms for perceptual grouping. Annual review of neuroscience 29, 203–227 (2006).

3. P. R. Roelfsema, R. Houtkamp, Incremental grouping of image elements in vision. Attention, Perception, and Psychophysics 73, 2542–2572 (2011).

4. K. Koffka, Principles of gestalt psychology. Principles of Gestalt Psychology 1–720 (2013). 10.4324/9781315009292/PRINCIPLES-GESTALT-PSYCHOLOGYKOFFKA.

5. M. Wertheimer, Untersuchungen zur Lehre von der Gestalt. II. Psychologische Forschung 4, 301–350 (1923).

6. J. R. Bergen, B. Julesz, Parallel versus serial processing in rapid pattern discrimination. Nature 303, 696–698 (1983).

7. P. Jolicoeur, S. Ullman, M. Mackay, Curve tracing: A possible basic operation in the perception of spatial relations. Memory & Cognition 14, 129–140 (1986).

8. M. W. Self, et al., The Effects of Context and Attention on Spiking Activity in Human Early Visual Cortex. PLOS Biology 14, e1002420 (2016).

9. A. Pooresmaeili, P. R. Roelfsema, A growth-cone model for the spread of object-based attention during contour grouping. Current Biology 24, 2869–2877 (2014).

10. H. S. Scholte, H. Spekreijse, P. R. Roelfsema, The spatial profile of visual attention in mental curve tracing. Vision Research 41, 2569–2580 (2001).

11. R. Houtkamp, H. Spekreijse, P. R. Roelfsema, A gradual spread of attention during mental curve tracing. Perception & psychophysics 65, 1136–1144 (2003).

12. P. A. McCormick, P. Jolicoeur, Predicting the shape of distance functions in curve tracing: Evidence for a zoom lens operator. Memory & Cognition 19, 469–486 (1991).

13. P. Jolicoeur, S. Ullman, M. Mackay, Visual Curve Tracing Properties. Journal of Experimental Psychology: Human Perception and Performance 17, 997–1022 (1991).

14. P. Jolicoeur, M. Ingleton, Size invariance in curve tracing. Mem Cognit 19, 21–36 (1991).

15. D. Jeurissen, M. W. Self, P. R. Roelfsema, Serial grouping of 2D-image regions with object-based attention in humans. eLife 5 (2016).

16. D. Domijan, M. Marić, A multi-scale neurodynamic implementation of incremental grouping. Vision Research 197, 108057 (2022).

17. S. Mollard, C. Wacongne, S. M. Bohte, P. R. Roelfsema, Recurrent neural networks that learn multi-step visual routines with reinforcement learning. PLOS Computational Biology 20, 1– 28 (2024).

18. L. Kirchberger, et al., The essential role of recurrent processing for figure-ground perception in mice. Science Advances 7, eabe1833 (2021).

19. T. Brosch, H. Neumann, P. R. Roelfsema, Reinforcement Learning of Linking and Tracing Contours in Recurrent Neural Networks. PLoS Computational Biology 11, e1004489 (2015).

20. H. Adeli, S. Ahn, N. Kriegeskorte, G. Zelinsky, Affinity-based Attention in Self-supervised Transformers Predicts Dynamics of Object Grouping in Humans. (2023).

21. T. Van Kerkoerle, M. W. Self, P. R. Roelfsema, Layer-specificity in the effects of attention and working memory on activity in primary visual cortex. Nature Communications 8, 1–14 (2017).

22. A. Pooresmaeili, J. Poort, A. Thiele, P. R. Roelfsema, Separable Codes for Attention and Luminance Contrast in the Primary Visual Cortex. Journal of Neuroscience 30, 12701–12711 (2010).

23. M. W. Self, T. van Kerkoerle, H. Supèr, P. R. Roelfsema, Distinct roles of the cortical layers of area V1 in figure-ground segregation. Current biology 23, 2121–2129 (2013).

24. J. D. Semedo, et al., Feedforward and feedback interactions between visual cortical areas use different population activity patterns. Nature Communications 13, 1–14 (2022).

25. D. Sridharan, E. I. Knudsen, Selective disinhibition: A unified neural mechanism for predictive and post hoc attentional selection. Vision Research 116, 194–209 (2015).

26. F. van der Velde, M. de Kamps, From Knowing What to Knowing Where: Modeling Object-Based Attention with Feedback Disinhibition of Activation. Journal of cognitive neuroscience 13, 479–491 (2001).

27. H. J. Pi, et al., Cortical interneurons that specialize in disinhibitory control. Nature 503, 521–524 (2013).

28. R. S. Sutton, A. G. Barto, Reinforcement Learning: An Introduction, Second Edition (2018).

29. D. P. Kingma, J. L. Ba, Adam: A Method for Stochastic Optimization. 3rd International Conference on Learning Representations, ICLR 2015 - Conference Track Proceedings (2014).

30. L. Kiorpes, S. A. Bassin, Development of contour integration in macaque monkeys. Visual Neuroscience 20, 567–575 (2003).

31. J. Atkinson, O. Braddick, Visual segmentation of oriented textures by infants. Behavioural Brain Research 49, 123–131 (1992).

32. A. Norcia, V. Sampath, C. Hou, F. Pei, Contour Integration in Human Infants. Investigative Ophthalmology & Visual Science 43, 3993–3993 (2002).

33. A. Needham, Infants’ use of featural information in the segregation of stationary objects. Infant Behavior and Development 21, 47–76 (1998).

34. D. M. Werchan, A. G. E. Collins, M. J. Frank, D. Amso, 8-Month-Old Infants Spontaneously Learn and Generalize Hierarchical Rules. Psychological Science 26, 805–815 (2015).

35. I. Pozzi, S. Bohte, P. Roelfsema, Attention-Gated Brain Propagation: How the brain can implement reward-based error backpropagation in Advances in Neural Information Processing Systems, (Curran Associates, Inc., 2020), pp. 2516–2526.

36. D. Linsley, A. Karkada Ashok, L. N. Govindarajan, R. Liu, T. Serre, Stable and expressive recurrent vision models in Advances in Neural Information Processing Systems, (Curran Associates, Inc., 2020), pp. 10456–10467.

37. W. Schultz, P. Dayan, P. R. Montague, A neural substrate of prediction and reward. Science 275, 1593–1599 (1997).

38. H. Roelfsema, P.R., Bohte, S. & Spekreijse, Algorithms for the Detection of Connectedness and Their Neural Implementation. Neuronal Information Processing: From Biological Data to Modelling and Applications. Series in Mathematical Biology and Medicine 7, 81–103 (1999).

39. I. Korjoukov, et al., The Time Course of Perceptual Grouping in Natural Scenes. Psychological Science 23, 1482–1489 (2012).

40. M. Ekman, P. R. Roelfsema, F. P. de Lange, Object selection by automatic spreading of top-down attentional signals in V1. The Journal of Neuroscience 40, 9250–9259 (2020).

41. T.-Y. Lin, et al., Microsoft COCO: Common Objects in Context. [Preprint] (2015).

42. B. A. Richards, et al., A deep learning framework for neuroscience. Nature Neuroscience 22, 1761–1770 (2019).

43. D. Schmid, H. Neumann, Thalamo-Cortical Interaction for Incremental Binding in Mental Contour-Tracing. bioRxiv 2023.12.20.572705 (2023). 10.1101/2023.12.20.572705.

44. D. Linsley, J. Kim, V. Veerabadran, C. Windolf, T. Serre, Learning long-range spatial dependencies with horizontal gated recurrent units in Advances in Neural Information Processing Systems, (Curran Associates, Inc., 2018).

45. L. Goetschalckx, et al., Computing a human-like reaction time metric from stable recurrent vision models in Advances in Neural Information Processing Systems, A. Oh, et al., Eds. (Curran Associates, Inc., 2023), pp. 14338–14365.

46. P. R. Roelfsema, F. P. de Lange, Early Visual Cortex as a Multiscale Cognitive Blackboard. Annual review of vision science 2, 131–151 (2016).

47. B. Hadad, D. Maurer, T. L. Lewis, The effects of spatial proximity and collinearity on contour integration in adults and children. Vision Research 50, 772–778 (2010).

48. S. Salehi, J. Lei, A. S. Benjamin, K.-R. Müller, K. P. Kording, Modeling Attention and Binding in the Brain through Bidirectional Recurrent Gating. [Preprint] (2024). Available at: https://www.biorxiv.org/content/10.1101/2024.09.09.612033v2.

49. S. P. Vecera, M. J. Farah, Is visual image segmentation a bottom-up or an interactive process? Percept Psychophys 59, 1280–1296 (1997).

50. J. Cohen, P. Cohen, S. G. West, L. S. Aiken, Applied multiple regression/correlation analysis for the behavioral sciences, 3rd ed. (Lawrence Erlbaum Associates Publishers, 2003).

